# Noisy models of the ventral stream reveal the impact of recurrence and learned representations on information processing timescales

**DOI:** 10.64898/2026.01.30.702784

**Authors:** Sara Varetti, Sebastian Goldt, Eugenio Piasini

## Abstract

In vision neuroscience, the temporal dynamics of the sensory stream and of its neural representations are thought to be deeply linked to the function of the hierarchy of cortical areas that deal with object recognition, known as the visual ventral stream. Neural representations that are invariant under identity-preserving object transformations, and therefore allow for efficient learning of object identity, are theorized to emerge from a self-supervised learning process that attempts to extract “temporally stable” features from the sensory input. Conversely, invariance increases along the hierarchy, putatively implying progressively slower codes in higher-level areas. Recent neurophysiological evidence shows that indeed, as one moves along this cortical hierarchy, neural representations of dynamic stimuli become slower, and additionally the temporal scales of the within-trial fluctuations of these representations (called “intrinsic timescales”) increase starkly. However, the network determinants of these timescale hierarchies are not understood in realistic systems, as the classical theory is based on models without noise, recurrence, or adaptive mechanisms. Here we investigate the temporal structure of the neural code in a noisy, recurrent and adaptive model of the ventral visual stream. We show that, surprisingly, the organization of the representation timescales is set by the broad architectural features of the network, regardless of training, while the intrinsic timescales depend on the details of the functions implemented on each layer. Our work underscores the importance of the temporal structure of the neural code as a probe for the link between structure and function in models of the vertebrate visual system.

## 1 Introduction

A major goal in visual neuroscience is to build models that capture the essential computational principles by which the visual cortex processes real-world inputs. In the last decade, convolutional neural networks (CNNs) have been found to be remarkably successful at modeling visual recognition from static images. This class of artificial neural networks has also demonstrated a remarkable ability to approximate cortical representations [1, 2], revealing hierarchical processing of inputs [3] and shedding light on the functional computations performed by the biological visual cortex [4, 5]. Despite their success in modeling cortical responses to static images, popular CNN-based models of visual cortex do not attempt to capture the inherently dynamic nature of visual experience, where information is continuously integrated across spatial [6, 7] and temporal [8] dimensions. After early attempts to use recurrence to improve the processing of static images [9] and theoretical insight that linked the depth advantage of certain residual networks to their mimicking of recurrent mechanisms [10], recently several architectures combining aspects of CNNs and recurrent networks (CNN-RNNs) have been proposed to bridge the gap between the spatial processing capabilities of convolutional networks and the need for temporal integration mechanisms. These architectures typically rely on recurrent connections to capture dynamic aspects of visual input [11–14]. Recurrent components are also included by large-scale efforts to build “foundation models” for visual neuroscience [15]. However, little attention has been devoted so far to the systematic study of the temporal properties of modeled activity. In particular, experimental evidence has shown that neural activity timescales are organized hierarchically, both within sensory cortex and across the brain more broadly, in humans, nonhuman primates, and rodents [8, 16–20]. An increase in the timescale, or the temporal stability, of neural codes is a crucial feature of classical and modern accounts of the emergence of invariance in sensory cortices [21–30]. Therefore, it is natural to ask under what conditions our models of visual cortex possess a similar property. To address this question, in this work we design and study a hybrid architecture that integrates convolutions, recurrent dynamics, adaptation and intrinsic noise, while preserving a simple structure that allows us to develop architectural parallels to visual cortex. The model is built on top of CORnet [14, 31] and from that architecture inherits a small number of layers that can be directly mapped onto areas of the ventral visual stream, enabling a biologically grounded investigation of temporal processing. We focus our investigation on two notions of timescales, as formalized by [19]: the timescale of the average neural representation of a stimulus (*response timescale*), and the timescale of random fluctuations around the average (*intrinsic timescale*). Our analyses show that, while a hierarchy of response timescales emerge robustly as a byproduct of the large-scale organization of network models, intrinsic timescales are sensitive to the fine details of a model and of its training. These results lead us to argue that intrinsic timescales are a useful conceptual construct that could guide the design and development of computational models of sensory systems.

## 2 Results

To quantify the timescales of neural codes for visual stimuli, we take inspiration from the empirical observations in [19], where electrophysiological recordings were made in rat visual cortex while the animal was exposed to dynamic visual stimuli (natural and synthetic videos) Fig 1. More specifically, data was collected from four cortical areas that constitute the rat analogue of the ventral visual stream — striate (V1), lateromedial (LM), laterointermediate (LI), and laterolateral (LL) visual cortex [32–34]. In [19], the temporal structure of the neural responses were characterized by two distinct measures: response timescales and intrinsic timescales. The response timescale captures the dynamics of the average neural response to a certain stimulus, and is expected to increase along the ventral stream by virtue of an increase in invariance of neural representations under identity-preserving transformations [26]. The intrinsic timescale depends only on the trial-by-trial fluctuations of the neural responses around the average (see *Estimation of Temporal Correlation* in Methods for more details); it is, in other words, the timescale of temporal noise correlations [18, 35, 36], and as such it has been proposed to be related to the role of recurrent and adaptive mechanisms in neural circuits [19]. Our goal in the present work is to identify the minimal set of mechanisms that a CNN-RNN architecture requires to reproduce this hierarchical organization of response and intrinsic timescales across cortical areas. To directly test which architectural elements are sufficient to reproduce these dynamics, we construct a family of CNN-RNN models capable of processing the same visual stimuli shown to the rats in the experiments. We build upon CORnet [14, 31] as a baseline model of the ventral stream and introduce a series of minimal but systematic modifications. We feed the movies to each model and measure the resulting timescales, yielding results that are directly comparable to those in [19] (note that CORnet is designed to model neural activity in primates, and not in rat, but as mentioned above the hierarchical organization of timescales across cortical areas is a general feature of neural activity shared by rodents and primates). A summary of the timescale dynamics resulting from each architectural modification is provided in Table 1. This stepwise approach allowed us to isolate the contribution of each component to the emergence of stable and temporally persistent activity patterns across the hierarchy of areas of the network.

**Table 1.**
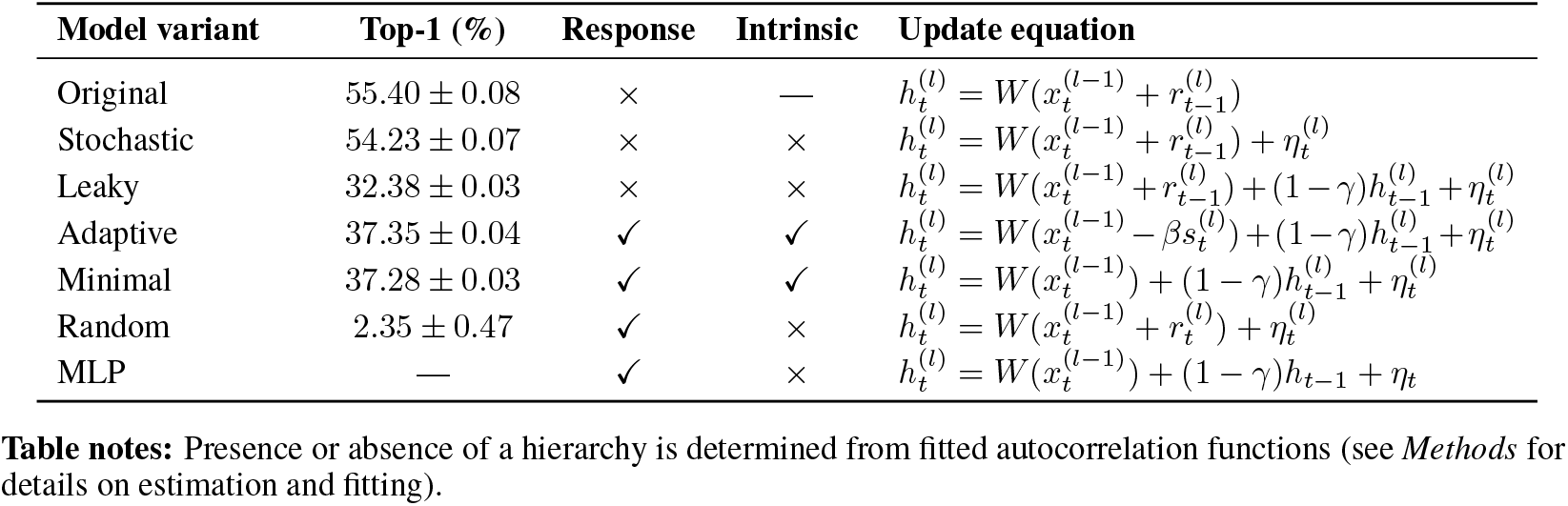
Summary of model variants and their effect on the emergence of a hierarchy of response and intrinsic timescales, as well as on image recognition performance. Each model builds incrementally on the original COR-net architecture with minimal modifications. Presence or absence of a temporal hierarchy is determined from fitted autocorrelation functions. Variables: 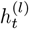 (hidden state at layer *l* at time *t*); 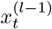 (input from previous layer); 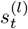 (adaptation variable); *γ* (leak); *β* (adaptation strength); 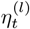 (noise).

**Figure 1.**
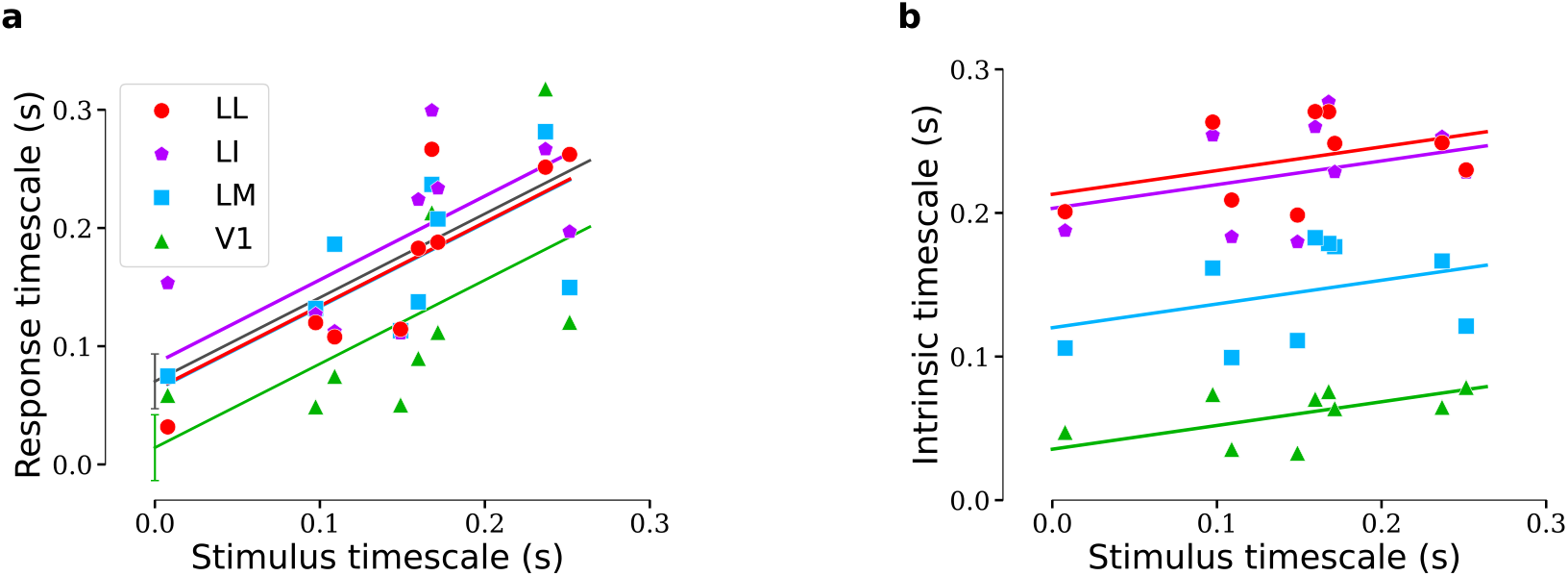
Response and intrinsic timescales in rat visual cortex, from [19]. Response a) and intrinsic timescales b) as a function of stimulus timescales for four visual areas (V1, LM, LI, LL). The stimulus timescale represents the decay time of the pixel-wise correlation between frames in each dynamic input (movie), i.e. how rapidly the visual content changes over time. Response, intrinsic, and stimulus timescales were estimated from neuronal recordings during presentation of naturalistic movie stimuli (see *Methods* for details on the computation of autocorrelation functions and the fitting procedure used to extract timescales).

### Stochastic CORnet

We begin by evaluating a biologically inspired architecture that combines the hierarchical structure of deep convolutional networks with local recurrent dynamics within each visual area. CORnet was designed as a minimal, cortex-mappable model of the ventral stream, with just four areas — V1, V2, V4, and IT — each performing canonical neural computations such as convolution, normalization, and ReLU nonlinearities. This simplicity stands in contrast to modern CNNs with multiple layers, making CORnet particularly suitable for comparisons with anatomical and functional properties of visual cortex [14, 31]. In CORnet, each area updates its state by integrating its current input with its own activity from the previous time step, according to:

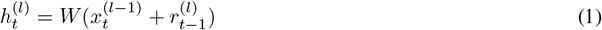

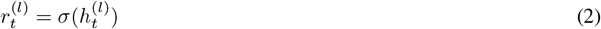

where 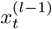 is the input from the preceding area, 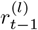 is the recurrent activation from the same area at the previous time step, and *W* represents the set of convolutions and normalizations. This recurrent update allows each layer to integrate information over time, a key feature missing in purely feedforward CNNs.

However, in this configuration, the model is entirely deterministic: its output is fully determined by the stimulus sequence. This makes it unsuitable for measuring intrinsic timescales, which require observing spontaneous fluctuations in neural activity across repeated presentations of the same input. As argued in [17], intrinsic timescales reflect the duration over which neural activity remains temporally correlated in the absence of stimulus variability—typically revealed only when internal variability (e.g., noise) is present. Therefore, while CORnet is a biologically plausible structure for exploring stimulus-driven dynamics and hierarchical response timescales, it lacks the internal variability necessary to investigate intrinsic dynamics. This motivates the introduction of stochastic noise as an additional component. We do so by adopting the following state update rule:

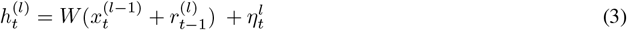

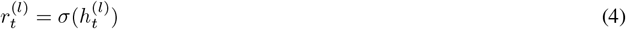

where 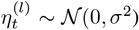 represents additive Gaussian noise injected into each layer to reproduce biological variability across trials (see Methods for details on the noise scaling).

We then tested whether this minimal model is capable of reproducing the experimentally observed hierarchy of response and intrinsic timescales. As mentioned above, this setup corresponds to a widely held hypothesis in computational neuroscience: that spatial invariance alone can lead to temporal stability. Under naturalistic stimulation, neurons in higher-order visual areas—such as IT—are thought to maintain more stable responses over time due to their invariance to object transformations, while early visual areas like V1 respond more transiently to low-level visual features rapidly changing in the input. Accordingly, one might expect that the progressive increase in spatial invariance along the ventral stream would be mirrored by a corresponding increase in temporal stability. CNNs, with their hierarchical architecture and increasing receptive field size, are designed to implement precisely such a progression.

However, our results show that spatial invariance alone is not sufficient to produce a biologically realistic hierarchy of timescales. Although response timescales show a weak increasing trend across areas, this pattern is stepwise rather than smoothly graded, and much less consistent than in the experimental data Fig 3. More strikingly, the intrinsic timescales show the opposite trend: they decrease along the hierarchy, Fig 3. In other words, activity in early visual areas like V1 remains temporally correlated over longer periods than in downstream regions, failing to reproduce the experimental findings from electrophysiological recordings.

**Figure 2.**
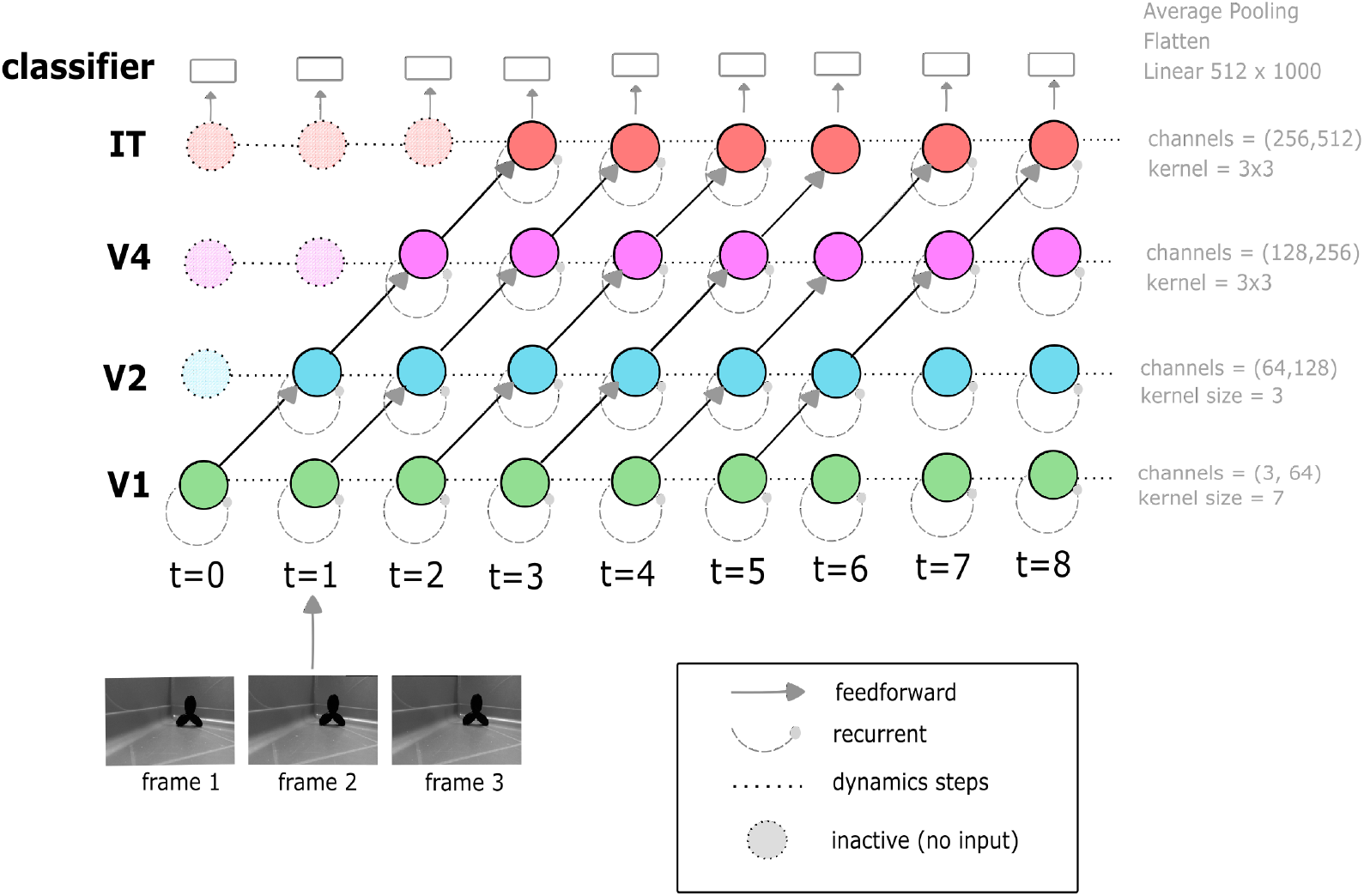
Network with recurrent and feedforward connections unrolled in biological time, introduced for static input in [14]. At each timestep, a new video frame is fed into V1, while the activity of higher visual areas (V2, V4, IT) evolves with a one-step delay relative to the previous layer. This staggered propagation causes information to flow sequentially through the hierarchy, so that at t=0 only V1 is driven by the input, at t=1 V2 becomes active, and so on, until the signal reaches IT after several timesteps. Dashed circles indicate the internal recurrent dynamics, while solid arrows mark the feedforward drive linking successive areas. This temporally unrolled scheme differs from conventional recurrent implementations, where all layers are updated simultaneously, and instead captures a more biologically realistic cascade of activation across the ventral stream.

**Figure 3.**
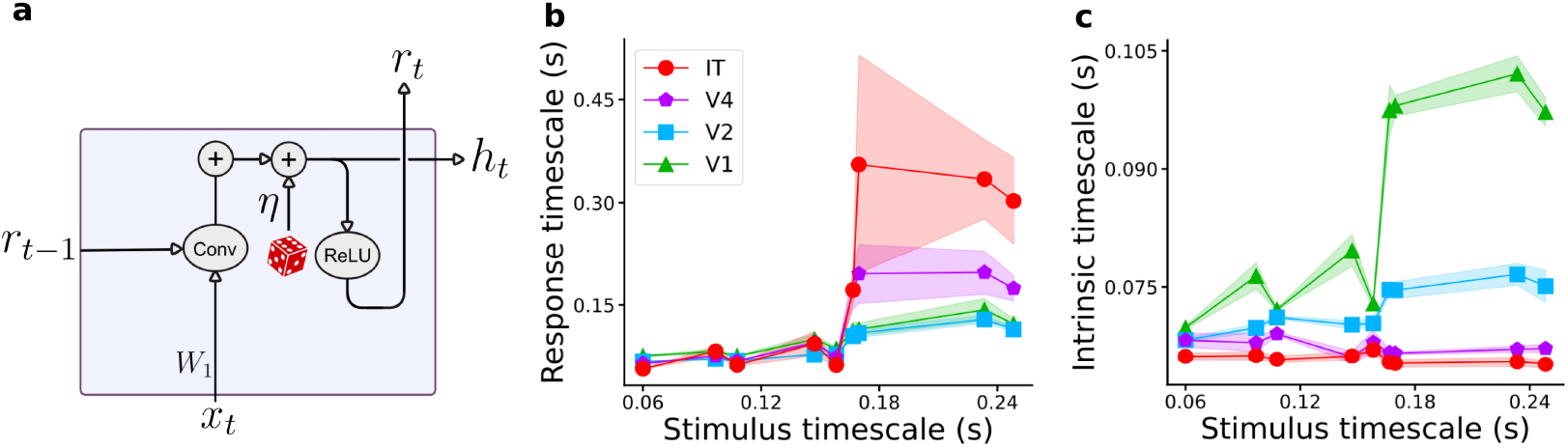
a) Circuitry within an area of the Stochastic CORnet model, corresponding to the equations Eq.4. Here, *h*_*t*_ is the pre-activation variable, *r*_*t*_ = *σ*(*h*_*t*_) is the post-activation unit activity (ReLU output), and *η*_*t*_ the additive Gaussian noise. *W*_1_ denotes the convolution that filters only the feedforward input *x*_*t*_ (from the visual stimulus or the preceding area), whereas the **Conv** block represents the second convolution integrating both the feedforward and the recurrent signals within the same area; b) Response timescales; c) Intrinsic timescales. For each area, the timescale was computed by averaging the estimates over 5 different subsets of 200 randomly selected units from the corresponding layer. Shaded areas represent the standard deviation across subsets (see Methods).

### Leaky CORnet

Increasing temporal stability can also arise from mechanisms beyond spatial invariance alone. A prominent alternative is the intrinsic dynamics of the network: neurons may exhibit a form of temporal inertia, whereby their current activity reflects a memory of recent past states, even when external inputs vary rapidly. This is in line by previous work suggesting that intrinsic timescales can emerge from autoregressive dynamics, where each state is shaped by its own temporal history [17, 18]. To test this possibility, we introduced a leak term in the network dynamics, controlling the auto-regressive character (or “inertia”) of neural activity. Specifically, for each area, the dynamics evolve according to:

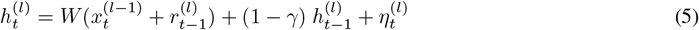

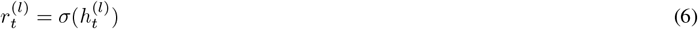

where *γ* modulates the balance between input-driven updates and persistence of previous states. Larger values of *γ* promote smoother temporal evolution of neural activity, even under dynamic stimuli, consistent with the autoregressive processes underlying intrinsic timescales. To model the hierarchical organization of temporal processing observed in the cortex, we assigned different values of *γ* to each area, decreasing along the visual hierarchy from V1 to IT. Specifically, we set *γ*_V1_ = 0.6, *γ*_V2_ = 0.5, *γ*_V4_ = 0.4, and *γ*_IT_ = 0.3. Since the effective leak rate is given by 1 − *γ*, areas with lower *γ* values retain more of their previous activity, resulting in a slower decay of autocorrelation over time.

However, the combination of input recurrence and leaky integration does not yield a consistent hierarchy of timescales across areas. Despite the presence of both feedback from prior responses and temporal inertia, this configuration fails to produce the progressive increase in timescales observed experimentally Fig 4.

**Figure 4.**
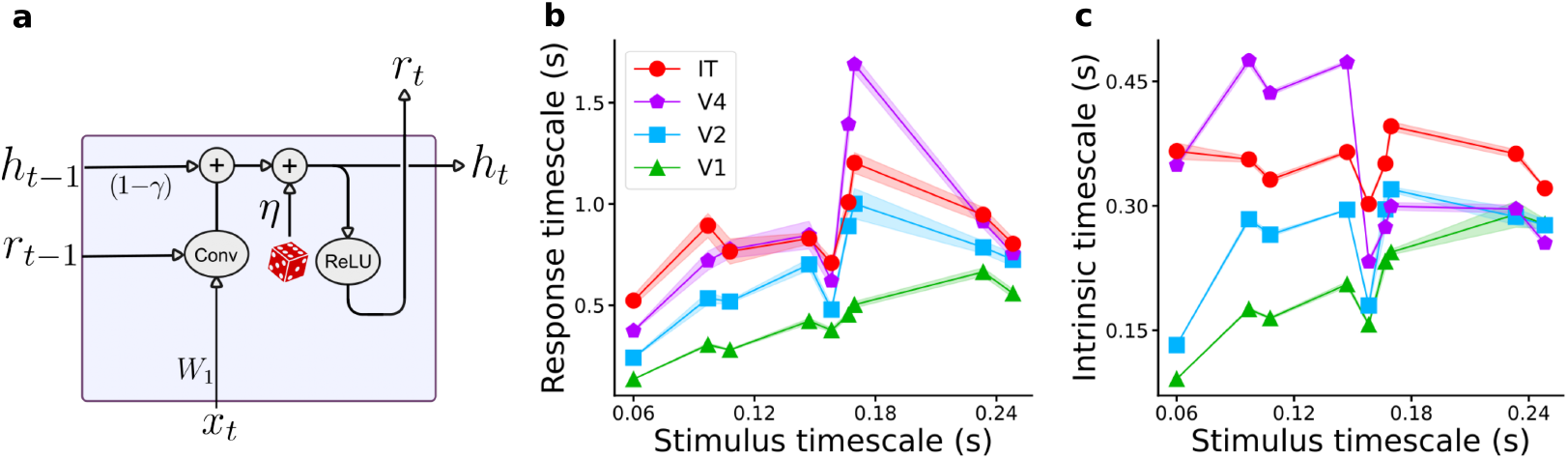
a) Circuitry within an area of the Leaky CORnet model, corresponding to the equations Eq.6. Relative to the Stochastic CORnet, this variant includes an effective leak rate 1 − *γ*, controlling the temporal integration of activity of the hidden state *h*_*t*_ over time; b) Response timescales; c) Intrinsic timescales. For each area, the timescale was computed by averaging the estimates over 5 different subsets of 200 randomly selected units from the corresponding layer. Shaded areas represent the standard deviation across subsets (see Methods).

### Adaptive CORnet

To further investigate the mechanisms that can shape intrinsic temporal dynamics beyond simple leaky integration, we introduced a model incorporating a form of neuronal adaptation. Specifically, we hypothesized that the current response of each neuron could be modulated by a memory trace of its past output, integrated over time with a tunable timescale. This idea is formalized through an auxiliary variable *s*_*t*_, defined as a low-pass filtered version of the past neuronal responses and modulate the input drive by subtracting this adaptation signal:

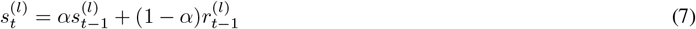

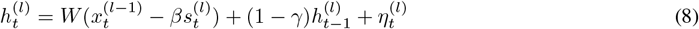

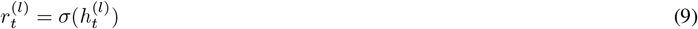

This formulation builds on the intrinsic suppression model by [37], augmenting it with noise and an autoregressive memory component. For each unit, *α* determines the integration window of the suppression mechanism. Larger *α* values correspond to longer memory and slower adaptation; smaller values emphasize recent activity. The other parameter, *β*, controls the strength of suppression. *β >* 0 yields activity-dependent suppression; *β <* 0 implements potentiation instead.

In analogy with the leaky model, the parameter *γ* was set independently for each area to reflect area-specific temporal integration. In contrast, *α* and *β* were kept constant across all layers [37].

We found that this model is able to reproduce the correct hierarchy of both response and intrinsic timescales Fig 5, at the same time allowing interactions among neurons within the same layer. Such cross-recurrence is essential in CNN-RNN models aiming to mimic cortical circuitry, as it provides the basis for population-level dynamics and spatially distributed temporal integration.

**Figure 5.**
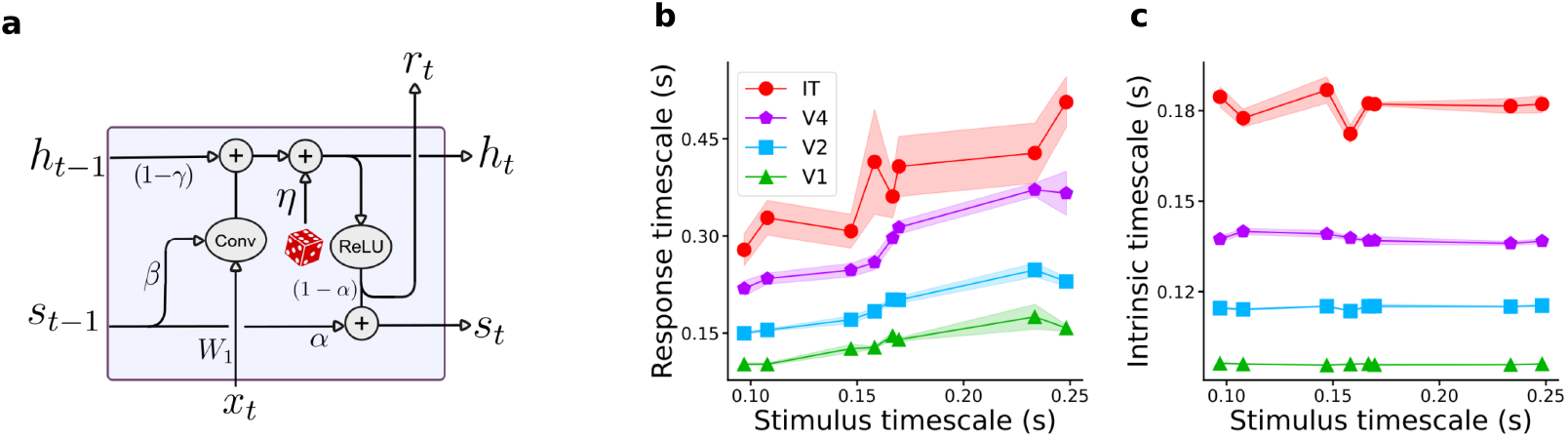
**a)** Circuitry within an area of the Adaptive CORnet model, corresponding to Eqs. 9. Relative to the Leaky CORnet, this variant introduces an adaptation variable *s*_*t*_, updated as a low-pass filtered version of the past neuronal activity with timescale *α*, and subtracted from the input drive with strength *β*. These two parameters jointly determine the temporal profile of adaptation, shaping how past activity suppresses or enhances current responses. b) Response timescales; c) Intrinsic timescales. For each area, the timescale was computed by averaging across 5 subsets of 200 randomly selected units. Shaded regions indicate the standard deviation across subsets (see Methods).

### A Minimal Model of Intrinsic Dynamics

Interestingly, we find that even a minimal modification of CORnet, introducing a stochastic drive and a simple leaky integration term, can give rise to a realistic hierarchy of timescales. Specifically, we consider the following simplified dynamics:

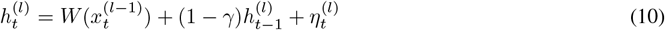

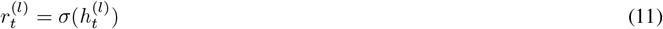

This formulation eliminates both recurrence from previous outputs and adaptation mechanisms, relying solely on the interaction between noisy inputs and leaky accumulation of past states. Despite its simplicity, this model captures a key feature of cortical dynamics: the progressive increase in temporal stability along the ventral stream. By assigning decreasing values of *γ* from early to late visual areas — thus increasing the effective memory of past activity — we obtain a monotonic hierarchy of intrinsic timescales that matches neurophysiological recordings Fig 6.

**Figure 6.**
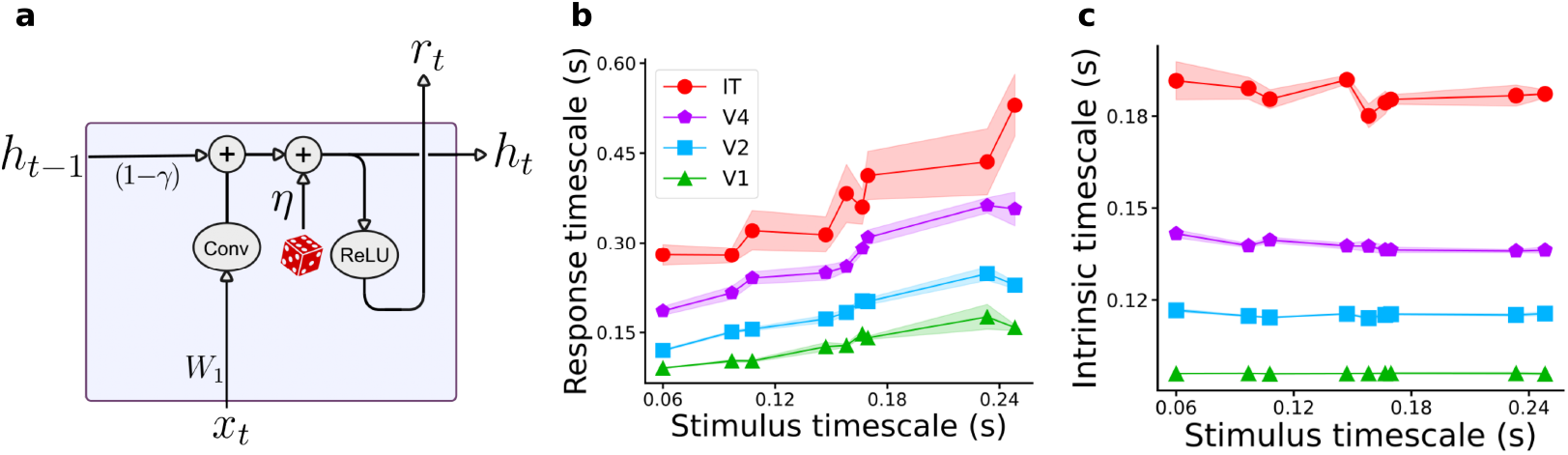
**a)** Diagram depicts circuitry within an area of Minimal CORnet, corresponding to Eq 11. Even without propagating the previous response *r*_*t*−1_ through the convolution *W*_1_ - neither as in the original CORnet nor in the adaptation model — and by introducing only an effective leak term (1 − *γ*), the network already exhibits a minimal form of intrinsic dynamics that reproduces the expected hierarchy of temporal processing across areas and relative to the stimulus timescale.; b) Response timescales; c) Intrinsic timescales. For each area, the timescale was computed by averaging the estimates over 5 different subsets of 200 randomly selected units from the corresponding layer. Shaded areas represent the standard deviation across subsets.

### The role of learned computations

To assess whether the emergence of temporal hierarchies depends on the learned computations of the network, we introduced a control version of the model in which all weights are randomly initialized rather than pretrained on ImageNet. This variant preserves the same dynamical architecture of the minimal model, i.e stochasticity and leaky integration, but removes any structured knowledge acquired through training. The motivation behind this manipulation is to disentangle the contributions of network architecture from those of learned representations. If the temporal hierarchy (i.e., increasing response timescales and intrinsic timescales across areas) were to persist in the absence of training, it would suggest that these properties emerge primarily from the network’s structural design. Conversely, if training is necessary for the correct hierarchy to appear, this would support the hypothesis that task-driven computations are essential for shaping the temporal dynamics observed in the ventral stream. Computational evidence suggests that although introducing a leak term and assigning area-specific *γ* values modulates the intrinsic persistence of neural activity, this mechanism alone is insufficient to generate a intrinsic temporal hierarchy. When the same leak dynamics are implemented in a network with randomly initialized weights — rather than using pretrained ImageNet weights — no systematic progression of intrinsic timescales across layers is observed Fig 7. This indicates that the temporal hierarchy observed in trained networks emerges from an interaction between intrinsic temporal integration and task-driven learning.

**Figure 7.**
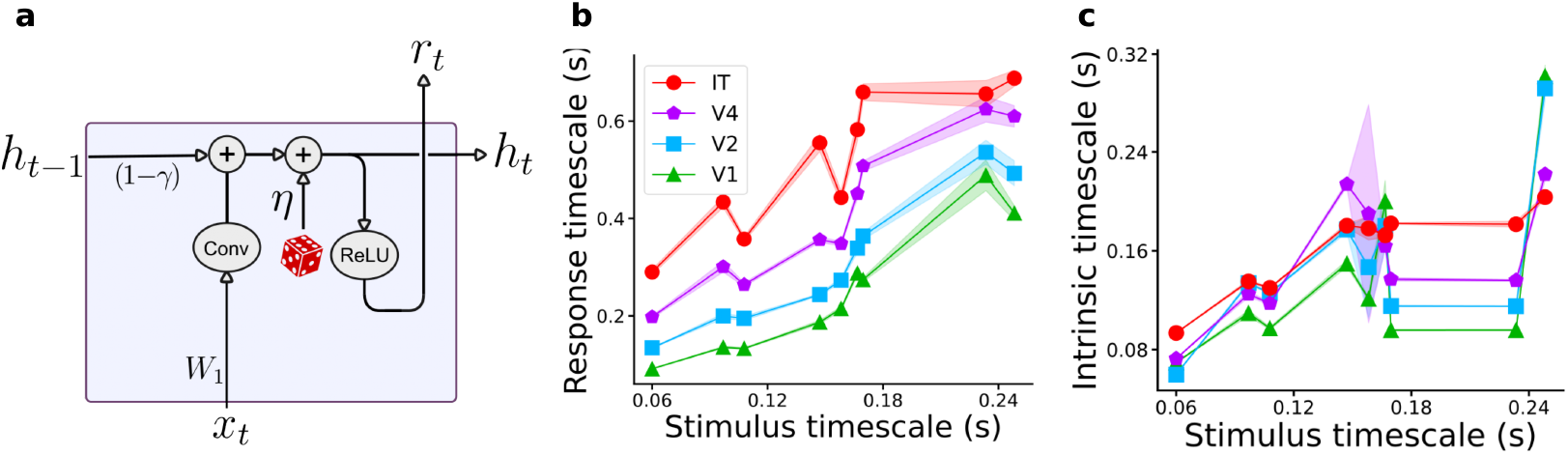
**a)** Diagram depicts circuitry within an area of Random CORnet. The architecture is identical to the Minimal CORnet model, except that all convolutional weights are randomly initialized rather than pretrained on ImageNet for object recognition; **b)** Response timescales; **c)** Intrinsic timescales. For each area, the timescale was computed by averaging the estimates over 5 different subsets of 200 randomly selected units from the corresponding layer. Shaded areas represent the standard deviation across subsets.

### Recurrent MLP: the effect of removing Convolutional Structure

To disentangle the effect of temporal recurrence *h*_*t−*1_ from architectural constraints such as convolutional structure, we designed a minimal recurrent multilayer perceptron (MLP) that processes each frame of a video as a flattened visual input. Each layer integrates its past activity via a leaky update rule, without spatially structured operations such as convolutions. This allows us to investigate the role of temporal recurrence alone, in the absence of any spatial hierarchy. Each layer implements a dynamics:

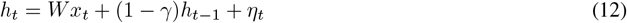

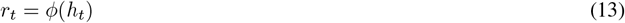

The matrix W represents a fixed random linear transformation mapping the input *x*_*t*_ to the hidden units. It is initialized once and kept constant. The network comprises four layers with fixed width Table 2. The MLP was (approximately) matched in terms of total number of parameters with CORnet-R; we note that in order to enforce this matching the input resolution for the MLP had to be significantly decreased (see Methods). Despite the lack of convolutional structure, the model still exhibits temporal dynamics via the diagonal recurrent connections within each layer, enabling a comparison with more complex CNN-RNN architectures.

**Table 2.** Architecture of the recurrent MLP model. Each layer implements leaky temporal integration with no spatial structure and processes the output of the previous layer, maintaining constant dimensionality from V2 onward.

| Layer | Input dimension | Number of units |
| --- | --- | --- |
| V1 | 3072 | 1024 |
| V2 | 1024 | 1024 |
| V4 | 1024 | 1024 |
| IT | 1024 | 1024 |

We compared the response and intrinsic timescales across layers of the recurrent MLP model with decreasing leak rate along the hierarchy to those of CORnet with random weights. As shown in Fig 8, deeper layers exhibit slower responses to dynamic inputs. This result highlights that diagonal recurrence alone — without any convolutional structure or learned weights — can produce a hierarchical organization of response timescales, that is, progressively slower stimulus-evoked activity across layers. However, this effect does not extend to intrinsic timescales which do not form a consistent hierarchy, mirroring what is observed in CORnet with random weights. These findings suggest that while response timescales can arise from simple architectural gradients such as varying leak rates, the emergence of intrinsic timescale hierarchies in network models may require more complex mechanisms, such as structured recurrent connectivity or learning-induced dynamics.

**Figure 8.**
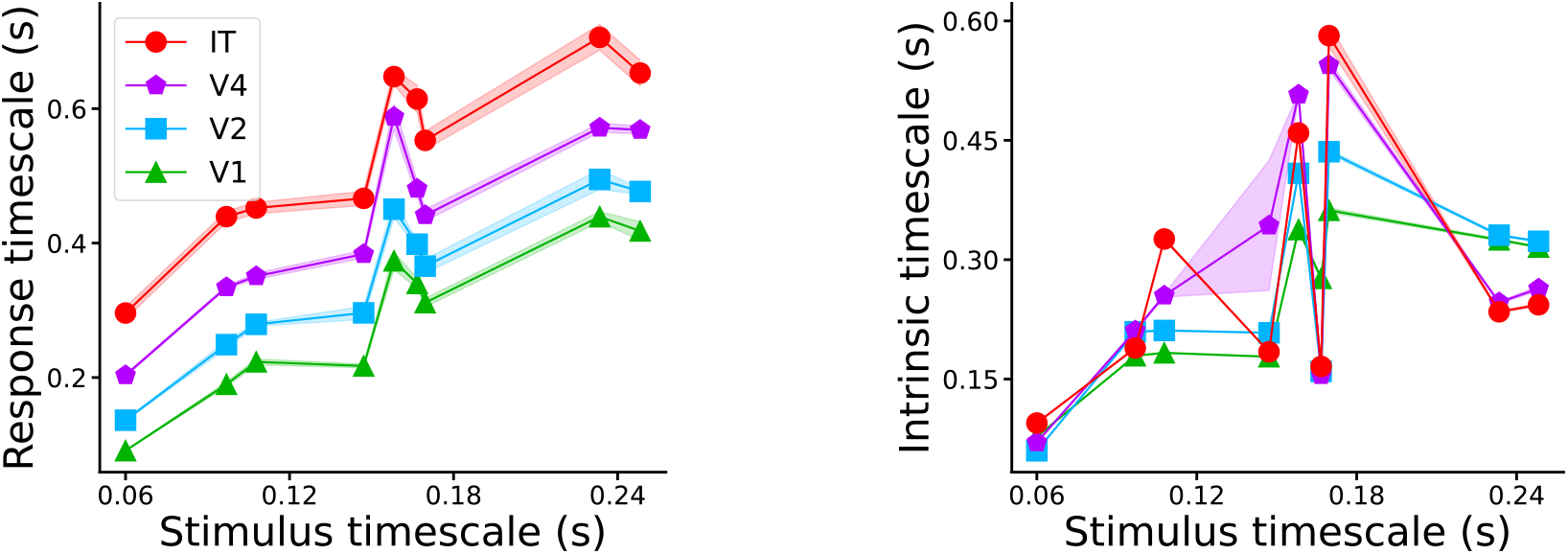
Response and intrinsic timescales in Randomly initialized chain of MLP. **a)** Response timescales; **b)** Intrinsic timescales. For each area, the timescale was computed by averaging the estimates over 5 different subsets of 200 randomly selected units from the corresponding layer. Shaded areas represent the standard deviation across subsets. Notably, intrinsic and response timescales exhibit lower variability compared to non-random cases, with average standard deviation across all points of ∼ 10^*−*3^

### ImageNet Classification Performance

To evaluate the image recognition capability of our modified CORnet-RT network in its cross-adaptation configuration, we adopted a standard linear probing protocol [38, 39]. We first extracted representations from the output layer of the IT block. For each image in the training and validation sets of ImageNet-1K, the model was run in evaluation mode for T = 5 recurrent time steps. This choice ensures consistency with the original CORnet training procedure. We retained the output of the final time step from IT, then applied average pooling followed by flattening to obtain a feature vector. These representations were then used to train a linear decoder. Feature extraction was performed in batches of 256 images. This approach, commonly used to assess representational quality, allows us to isolate the contribution of the learned features independently of the classifier training, and to test whether the internal modifications introduced in our model architecture preserve linearly decodable object representations. Despite no end-to-end fine-tuning, the minimal model yields a top-1 accuracy of 37.28% and a top-5 accuracy of 62%, indicating that the internal recurrent dynamics preserve linearly decodable high-level object representations. We note this result outperforms the accuracy reported for the recent CordsNet-R4 model (33.78%) [12], despite CordsNet undergoing an elaborate three-stage initialization and fine-tuning procedure specifically designed to optimize performance in continuous-time RNNs. The fact that it achieves competitive performance with significantly simpler training highlights the computational efficiency and architectural robustness of our design. To estimate the variability of linear decoding performance, we trained the linear classifier multiple times (N=3) with different random seeds, while keeping the extracted representations fixed.

For reference, the top-1 and top-5 validation accuracies of the original CORnet-RT models tested on ImageNet-1K are: 55.4%top-1, 78.9% top-5. In the following, we report the classification performance of all the model variants introduced to explore the hierarchy of timescales Table 3. The accuracy progression of the linear decoder across training epochs is reported in Supplementary Figure 1.

**Table 3.** Top-1 and Top-5 classification accuracy for all model variants on the ImageNet-1K validation set. Accuracy values are reported as mean ± standard deviation across multiple validation runs.

| Model | Top-1 Accuracy (%) | Top-5 Accuracy (%) |
| --- | --- | --- |
| CORnet-RT | 55.40 ± 0.08 | 78.90 ± 0.03 |
| Stochastic CORnet-RT | 54.23 ± 0.07 | 78.17 ± 0.03 |
| Leaky CORnet-RT | 32.38 ± 0.04 | 56.30 ± 0.04 |
| Adaptive CORnet-RT | 37.35 ± 0.04 | 61.60 ± 0.02 |
| Minimal CORnet-RT | 37.28 ± 0.03 | 61.55 ± 0.02 |
| Random CORnet-RT | 2.35 ± 0.47 | 6.75 ± 1.13 |

## 3 Discussion

Our results demonstrate that the emergence of hierarchical temporal dynamics in convolutional recurrent networks critically depends on the interaction between architectural features and dynamical mechanisms. In particular, while response timescales increase along the hierarchy even in untrained networks, intrinsic timescales only exhibit the experimentally observed progression when explicit memory mechanisms (such as leaky integration) and learned representations are in place. First, we showed that spatial invariance alone, as implemented in the Noisy CORnet-R model, is not sufficient to account for the hierarchy of intrinsic timescales observed in cortical recordings. This directly challenges the hypothesis that temporal stability of responses necessarily follows from increasing spatial pooling and feedforward depth. Second, introducing leaky integration in the Leaky-CORnet model successfully recovered the intrinsic timescale hierarchy, confirming that slow internal dynamics — controlled here by area-specific leak parameters — are sufficient to produce long-lasting activity fluctuations. This result aligns with prior experimental findings linking intrinsic timescales to local autoregressive dynamics in cortical circuits [17, 18]. Third, adding cross-recurrence through adaptation mechanisms did not disrupt the hierarchy, as long as intrinsic memory was retained. Interestingly, we found that models in which the adaptation term dominated (i.e., with strong *β*) were prone to pathological auto-correlation profiles, limiting the biological plausibility of pure adaptation-based accounts. Instead, the combination of leaky dynamics and moderate adaptation yielded robust temporal hierarchies, supporting the idea that multiple recurrent mechanisms may coexist in shaping cortical dynamics. Finally, we found that training plays an essential role: in randomly initialized networks, the intrinsic hierarchy collapsed even when the architectural and dynamical features were preserved. This indicates that learned feature selectivity interacts nontrivially with internal dynamics, supporting the idea that intrinsic timescales are shaped by both structure and function. This complements and extends previous observations made in trained RNNs for action recognition tasks [40], and highlights the role of training even in models designed to reflect cortical connectivity. Taken together, these findings point to a multifactorial origin of temporal hierarchies in visual cortex, involving both structural motifs (such as hierarchical depth and local recurrence) and functional constraints (such as task-driven learning and internal memory). In our view, evaluating models of cortical computation should go beyond static object recognition and include a dynamical characterization of both evoked and intrinsic timescales. A key limitation of our current approach is that the model is not trained end-to-end on an object recognition task. Instead, it relies on fixed pretrained weights obtained from CORnet-RT trained on ImageNet. While this allows us to isolate the effect of architectural and dynamical modifications on temporal processing, it likely underestimates the full representational capacity of the model. In particular, we expect that retraining the modified architecture end-to-end could improve classification performance, potentially matching or exceeding the accuracy levels of the original CORnet. Future work should explore whether such end-to-end training affects the emergence and structure of temporal hierarchies, and whether performance gains align with changes in intrinsic or response timescales. CORnet was originally designed as a model of the primate ventral visual cortex, whereas our comparisons are based on experimental data recorded from the rat visual system. Nevertheless, the dynamical principles examined here — such as leaky integration, adaptation, and recurrence — are likely conserved across species, reflecting general strategies for balancing temporal integration and responsiveness [41–45].

Our framework opens several directions for future work. One important extension would be to investigate the role of feedback connections in shaping temporal hierarchies. In biological circuits, feedback interacts continuously with sensory input, modulating integration and response timescales across areas. Incorporating feedback pathways that operate on comparable timescales to the feedforward drive could therefore reveal new forms of dynamic across the cortical hierarchy. Another promising direction would be to decouple the intrinsic dynamical timescale of the model from the frame rate of the visual input. In our current implementation, the temporal update of neural activity coincides with the presentation of consecutive video frames, effectively constraining the system’s dynamics to the timescale of the stimulus. Allowing the network to evolve on a finer temporal resolution — or to integrate multiple internal steps per frame — could uncover richer temporal behaviors, including oscillatory or predictive regimes that may otherwise be suppressed by forcing the dynamics to unfold in lockstep with the movie frames.

## 4 Methods

### Network Architecture

CNN models operate in a purely feedforward manner, lacking the rich lateral connections, and the complex response dynamics they produce, that are known to characterize the visual flow of the ventral region. While classical CNNs trained for image recognition have achieved remarkable success, they remain incomplete models of the visual system, as they lack mechanisms for temporal integration. To address this limitation, the CORnet family of models [14] has been proposed as a more biologically inspired architecture for visual processing. In particular, CORnet-R introduces local recurrent dynamics within each visual area (V1, V2, V4, IT), while maintaining a hierarchical feedforward structure. Each area performs biologically plausible computations, including convolutions, normalization, ReLU nonlinearities, and pooling (see Table 4 for details). The architecture is unrolled in time in a biologically realistic manner: At time *t* = 0, only the first layer is active; the second layer receives input from the first one at *t* = 1, etc, resulting in an input reaching the last layer only after four steps. Unlike standard machine learning implementations of recurrent models, which propagate input through all layers simultaneously at each time step, this stepwise temporal unrolling mimics the biologically plausible flow of information across areas Fig 2. As such, it provides a useful framework for investigating hierarchical neural dynamics. The standard CORnet model ends with a linear decoder trained on ImageNet. At each time step, activity from the final stage is flattened and passed to a classifier, enabling object recognition across 1000 categories via a softmax layer. In the original CORnet-RT model, each visual area evolves over a fixed number of discrete time steps (typically times = 5), but the input image remains static throughout: at *t* = 0, the image is presented to V1, which updates its internal state; at *t* = 1, V1 passes its output to V2, and so on. This leads to temporal dynamics across areas, but they are not driven by a time-varying stimulus. To model neural responses to dynamic input, we modified CORnet-RT to process a sequence of video frames, feeding a new frame to V1 at each time step. Unlike the original implementation - which retains only the final output - we also store the full time sequence of activations across all areas. This enables detailed analysis of the evolving internal dynamics and allows us to compare model predictions to neural recordings obtained with movie stimuli. To mimic biological trial to trial variability, we introduced, in the second convolutional layer of each area, gaussian noise *η*_*t*_ ∼ 𝒩 (0, *σ*^2^) where *σ* = noise_area_ · *α* with noise_area_ = 0.7 for each area and *α* = 10^*−*5^. The scaling factor *α* provides an additional level of control over the noise magnitude, allowing for experimentation with different noise intensities. We selected the minimum value of *α* that preserved the model’s functional performance while introducing sufficient variability across trials to estimate intrinsic timescales. As shown in Table 5, *α* = 1 × 10^*−*5^ leads to only a minimal drop in classification accuracy relative to the original CORnet-RT, maintaining a top-1 accuracy of 54.23% and a top-5 accuracy of 78.17% on ImageNet-1K, compared to 55.4% and 78.9% respectively. Higher noise values severely compromise classification performance. All model variants share the same architectural backbone, temporal unfolding mechanism, and stochastic drive described above. To explore the role of different neural mechanisms in shaping intrinsic dynamics, we manipulated the recurrent update equations by including additional components—namely, leaky integration and adaptation. These components were instantiated by tuning a small set of hyperparameters (*γ, α, β*), whose values were kept fixed across trials and stimuli. The table 6 summarizes the parameter values used in each variant.

**Table 4.** Architectural parameters of the CORnet model. Each stage corresponds to one cortical area (V1, V2, V4, IT). The decoder stage performs average pooling, flattening, and a final linear layer producing 1000-class outputs.

| Stage | Operation | Output shape |
| --- | --- | --- |
| Input | – | $224 \times 224 \times 3$ |
| V1 | conv $7 \times 7/4$ | $56 \times 56 \times 64$ |
| | conv $3 \times 3$ | $56 \times 56 \times 64$ |
| V2 | conv $3 \times 3/2$ | $28 \times 28 \times 128$ |
| | conv $3 \times 3$ | $28 \times 28 \times 128$ |
| V4 | conv $3 \times 3/2$ | $14 \times 14 \times 256$ |
| | conv $3 \times 3$ | $14 \times 14 \times 256$ |
| IT | conv $3 \times 3/2$ | $7 \times 7 \times 512$ |
| | conv $3 \times 3$ | $7 \times 7 \times 512$ |
| Decoder | avgpool | $1 \times 1 \times 512$ |
|  | flatten | 512 |
|  | linear | 1000 |

**Table 5.** Top-5 accuracy of the Stochastic CORnet-R model as a function of noise magnitude *α*.

| Noise scale $\alpha$ | Top-5 Accuracy (%) |
| --- | --- |
| 0.0001 | 78.45 |
| 0.0002 | 74.52 |
| 0.0003 | 41.89 |
| 0.0004 | 10.40 |

Increasing the noise magnitude $\alpha$ progressively degrades the model’s classification performance, with accuracy dropping sharply beyond $\alpha = 0.0002$ .

**Table 6.** Model parameters for different CORnet variants. The values of *γ* control leaky integration in each area, while *α* and *β* define the adaptation strength and memory. Dashes indicate that the corresponding mechanism was not used. Each CORnet-R variant uses distinct dynamical parameters.

| Model variant | $\gamma_{V1}$ | $\gamma_{V2}$ | $\gamma_{V4}$ | $\gamma_{IT}$ | $\alpha$ | $\beta$ |
| --- | --- | --- | --- | --- | --- | --- |
| Leaky CORnet-R | 0.6 | 0.5 | 0.4 | 0.3 | – | – |
| Adaptive CORnet-R | 0.6 | 0.5 | 0.4 | 0.3 | 0.96 | 0.1 |
| Minimal CORnet-R | 0.6 | 0.5 | 0.4 | 0.3 | – | – |

### Stimuli and Network Presentation Protocol

The main stimulus set comprised nine video clips, each lasting 20 seconds and sampled at 30 frames per second (fps). The stimuli are described in depth in [19]. Briefly, six of the videos were naturalistic movies depicting real-world dynamic scenes, while the remaining three were synthetic controls: phase-scrambled versions of two natural movies and a white noise movie. The phase-scrambled movies were obtained by performing a spatiotemporal fast Fourier transform (FFT) over the standardized 3D array of pixel values, obtained by stacking the consecutive frames of a movie. The white-noise movie was generated by randomly setting each pixel in each frame either to white or black. Each video was composed by a sequence of 600 frames (20 s × 30 fps), and each frame was input to the models we analyzed at a single time step t, such that the recurrence in the network (i.e., the update of internal states *h*_*t*_) coincided with the temporal evolution of the movie. In this formulation, each change in frame corresponds to a single recurrence of the internal dynamics governed by the update rule:

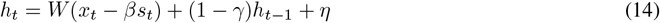

### Estimation of Temporal Correlation

To quantify the temporal structure of both stimulus and neural signals, we computed the characteristic timescale via the lagged Pearson correlation function:

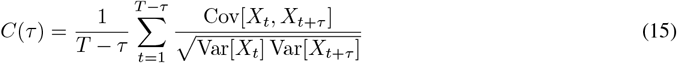

This metric captures how quickly a time series decorrelates over time and was applied uniformly across three domains of analysis:

1. **Movie stimuli**: *X*_*t*_ represents the pixel intensity vector of frame t. For each lag *τ*, we computed the average correlation coefficient between all frame pairs separated by *τ*, yielding a global measure of how rapidly the visual content changed over time. This allowed us to assign a characteristic timescale to each movie.
2. **Stimulus-driven neural activity**: *X*_*t*_ corresponds to the population vector of trial-averaged and temporally centered spike counts across neurons at time *t*, i.e., *X*_*t*_ = {*x*_*n*_(*t*) − ⟨*x*_*n*_⟩ _*t*_}. This approach mirrors the one used for the movies and characterizes the decay of evoked population activity across time lags.
3. **Intrinsic neuronal activity**: For each neuron, *X*_*t*_ was defined as the vector of activities across trials at time bin t, i.e., *X*_*t*_ = {*x*_*k*_(*t*)}, with k indexing trials. The autocorrelation *C*(*τ*) was computed for each unit and then averaged across all neurons within a given area before extracting a characteristic time constant, following [17] and [19].

This unified correlation framework enables direct comparisons of temporal scales across visual input and neural responses, both evoked and intrinsic.

To estimate the characteristic timescale from these autocorrelation functions, following [19] we fit the empirical curves using a damped oscillatory model:

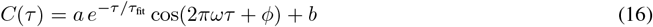

Here, *a* is the amplitude, *b* the long-term offset, *τ*_fit_ the decay timescale, *ω* the oscillation frequency, and *ϕ* a phase shift. Parameters *θ* = (*a, b, τ*_fit_, *ω, ϕ*) were optimized by minimizing the squared error:

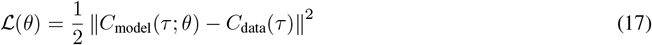

To avoid sensitivity to local minima, we used the differential evolution algorithm implemented in SciPy, with conservative parameter bounds, a population size of 250, and up to 500 iterations.

This fitting procedure was applied independently to each movie and to each cortical area (V1, V2, V4, IT). The fitted decay constant *τ*_fit_ served as a quantitative index of the timescale for both visual and neuronal autocorrelations. Model fits were visually inspected by overlaying the empirical and fitted curves.

### Design of the Multilayer Perceptron Architecture

Our choice of parameters was guided by the need to match, as closely as possible, the total number of parameters of the CORnet-R architecture (approximately 4.7 million). To achieve this, we treated the parameter count as a design constraint: because each layer of the MLP performs a fully connected transformation rather than a convolutional one, every unit receives input from the entire visual field. As a result, maintaining the original input resolution would have led to an unfeasibly large number of parameters. We therefore reduced the input dimensionality to 3 × 32 × 32, which allows the overall parameter count to remain comparable to that of CORnet-R (around 6 million in our implementation) while preserving access to global spatial information in each layer.

### Evaluation of representational quality

To evaluate the representational quality of CORnet in its modified configurations, we employed a standard linear probing approach. All experiments were conducted in PyTorch using standard ImageNet preprocessing and evaluation protocols. Representations were extracted from the decoder output after five recurrent time steps, and a linear classifier was trained on frozen features. The full set of implementation parameters is reported in Table 7.

**Table 7.** Implementation details for the linear probing protocol used to evaluate CORnet-RT representations.

| Component | Details |
| --- | --- |
| Framework | PyTorch 1.11.0 |
| Data | ImageNet 2012 (Russakovsky et al., 2015) |
| Preprocessing | Resize to 256, center crop to $224 \times 224$ px, normalization using ImageNet mean and standard deviation |
| Representations | Extracted from the CORnet-RT decoder (flatten layer) with 5 recurrent time steps |
| Train set size | 1.28M images (ImageNet training split) |
| Validation set | 50K images (ImageNet validation split) |
| Classifier | Linear layer (input dimension = 512, output dimension = 1000) |
| Training | Features frozen; only the linear layer is trained |
| Loss | Cross-entropy |
| Optimizer | Adam |
| Learning rate | $1 \times 10^{-3}$ |
| Batch size | 256 |
| Epochs | 1 (per run; repeated with different random seeds) |
| Metrics | Top-1 and Top-5 accuracy (validation set) |

## Data and code availability

All stimuli (naturalistic movies, phase-scrambled controls, and white noise sequences) used in this study are publicly available at: https://osf.io/7gteq/. The code used to generate the results and figures in this paper will be made available upon request.

## Supplementary Material

### Accuracy over Epochs for the Linear Decoder

**Supplementary Figure 1.**
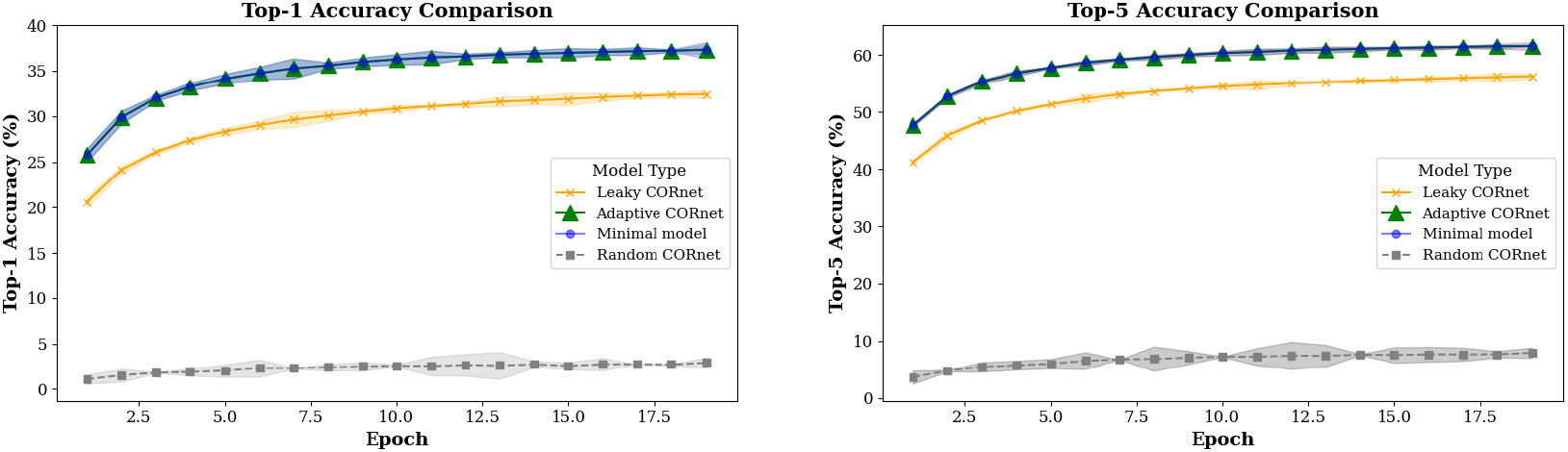
Linear decoding accuracy for pretrained CORnet-RT representations. (Left) Top-1 accuracy. (Right) Top-5 accuracy. A linear decoder was trained on frozen CORnet-RT features extracted from pretrained weights. Values represent the average over three independent training runs. Error bars (standard deviation) are scaled by a factor of 10 for improved visualization.

## Acknowledgments

We would like to thank Nawal Yahiaoui for contributing to the initial stages of the project.

SG gratefully acknowledges funding from the European Research Council (ERC) for the project “beyond2”, ID 101166056; from the European Union–NextGenerationEU, in the framework of the PRIN Project SELF-MADE (code 2022E3WYTY – CUP G53D23000780001), and from Next Generation EU, in the context of the National Recovery and Resilience Plan, Investment PE1 – Project FAIR “Future Artificial Intelligence Research” (CUP G53C22000440006). EP acknowledges funding from the European Union–NextGenerationEU, in the framework of the PRIN Project no. 2022XE8X9E - CUP G53D23004590001.

